# Graph-KIR: Graph-based KIR Copy Number Estimation and Allele Calling Using Short-read Sequencing Data

**DOI:** 10.1101/2023.11.29.568665

**Authors:** Hong-Ye Lin, Hui-Wen Chuang, Tsung-Kai Hung, Ting-Jian Wang, Ching-Jim Lin, Jacob Shujui Hsu, Chia-Lang Hsu, Ya-Chien Yang, Pei-Lung Chen, Chien-Yu Chen

## Abstract

**Motivation:** The Killer-cell Immunoglobulin-like Receptor (KIR) is a highly polymorphic region in the human genome, associated with autoimmune diseases and organ transplantation. The sequences of KIR genes are highly similar among star alleles as well as in between individual genes, with the copy number of each KIR gene typically ranging from 0 to 4. In this study, we introduce a tool Graph-KIR that aims to estimate the copy number of genes and to call full-resolution (7-digit) KIR alleles from a whole genome sequencing (WGS) sample.

**Results:** Graph-KIR, unlike most KIR tools, is capable of independently typing KIR alleles per sample with no reliance on the distribution of any framework gene in a cohort. In a set of 100 simulated samples, Graph-KIR demonstrated 100% accuracy in copy number estimation and high accuracy of allele typing: 91.2% at 7-digit resolution, 97.0% at 5-digit resolution, 97.2% at 3-digit resolution, and 99.6% at gene-level resolution. Graph-KIR outperforms existing tools such as PING’s WGS version (91.9% accuracy) and T1K (84.6% accuracy) at 5-digit resolution. By analyzing the results on 44 HPRC samples, Graph-KIR achieves an accuracy of 85.0%, better than PING’s WGS version (75.5% accuracy) at 5-digit resolution. The release of Graph-KIR adds another valuable tool to assist users in accurately estimating copy numbers and calling alleles of KIR genes from WGS samples, ensuring reliable performance.

**Availability:** The Graph-KIR and paper-related pipeline codes are available at https://github.com/linnil1/KIR_graph.

## 1 Introduction

The Killer-cell Immunoglobulin-like Receptor (KIR) family plays a crucial role in the operation of natural killer (NK) cells, controlling the cell-killing mechanism through its binding with Major Histocompatibility Complex (MHC) class I molecules on the cell surface [1]. Such interactions are gene-specific to both the Human Leukocyte Antigen (HLA) and KIR. Some studies [2, 3] have shown that certain ligations are gene content-specific, meaning that they depend not only on the types of genes present but also on the star alleles. Consequently, accurately calling the star alleles of all KIR genes in an individual is of significant importance for future research and clinical applications.

The KIR gene family, ranging from 70 to 270 kilobase pairs (kbp) in length, is located on chromosome 19 (19q13.4). This family encompasses 15 genes (*KIR2DL1/L2/L3/L4, KIR2DL5A, KIR2DL5B, KIR2DS1/S2/S3/S4/S5, KIR3DL1/S1, KIR3DL2/L3*) and two pseudogenes (*KIR2DP1* and *KIR3DP1*). All KIR genes, spanning approximately 5 to 13 kbp each, exhibit a notable similarity in both their DNA sequence structures and contents. Aside from the similarity across genes, the complex internal similarity within each gene is akin to another immune gene family, HLA. The star allele of a KIR gene, for instance, *KIR2DL1*0010101*, signifies a set of single nucleotide polymorphisms (SNPs) across the entire gene. The classification of star alleles is divided into three distinct resolution levels: identical protein sequences, identical DNA exon regions, and identical DNA sequences, corresponding to 3-digit, 5-digit, and 7-digit, respectively, according to the KIR nomenclature definition [4]. Each KIR gene typically has 0 to 4 star alleles in an individual, based on the KIR haplotype-level polymorphism, the value we refer to as the copy number (CN). All of these concepts are illustrated in Figure 1.

**Figure 1.**
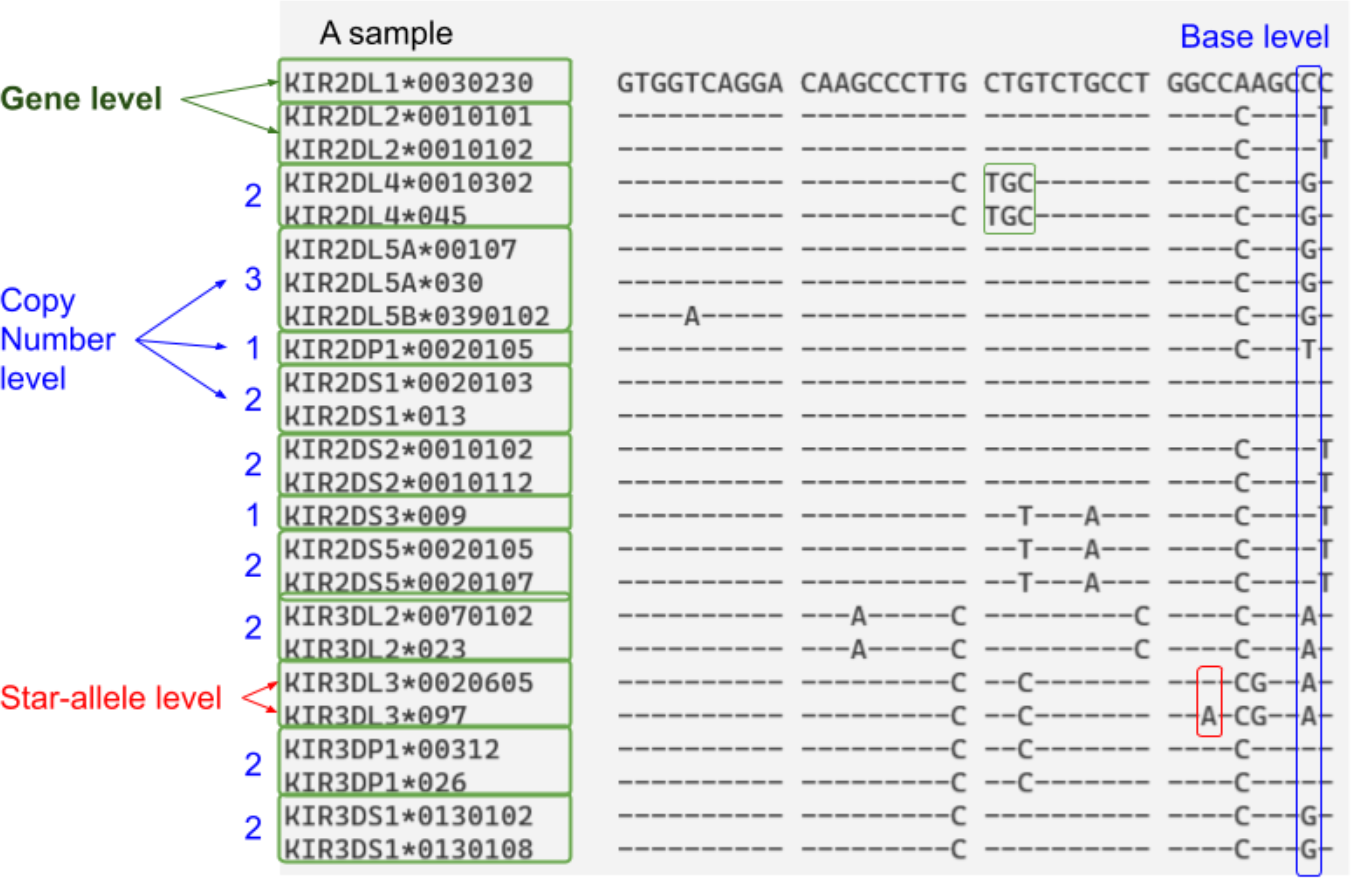
Demonstration of KIR complexity at four levels using one of the simulated samples. These levels comprise the base level, gene level, star-allele level, and copy number level, with the copy number level highlighting the count of star alleles within a single KIR gene. The symbol ‘-’ stands for a position identical to the base of *KIR2DL1*\*0030230.

KIR typing is usually divided into two tasks: Copy Number Estimation (addressing the gene level and copy number level) and Allele Typing (dealing with the star-allele (SNP) level). These two steps are crucial for KIR typing and are the common components of existing KIR tools. To our knowledge, there are five tools capable of identifying KIR genes from high-throughput genotyping or sequencing data: KIR*IMP [5], KPI [6], PING [7, 8, 9], the pipeline (denoted as SakaueKIR in this study) developed by Sakaue et al. [10], and T1K [11]. However, KIR*IMP does not support short-read sequencing data and KPI lacks the allele typing step. In a strict sense, only three tools are capable of accepting read files in FASTQ format as input to generate KIR allele typing results.

For the task of copy number estimation, one typical way is to determine the copy numbers of genes based on their corresponding read depths. The CN of a gene is highly related to its read depth in the short-read sequencing data. For example, when there are three alleles belonging to *KIR2DL1* (CN=3), all the reads originating from those three alleles will be mapped to *KIR2DL1*, resulting in the read depth of *KIR2DL1* being about 1.5 times higher than the normal read depth (i.e., diploid, CN=2). By gathering the read depths of genes, those values can be clustered into copy number groups by specific thresholds. PING carefully selects numerous probes to determine the read depths of each gene by counting the reads matching the probes followed by normalization based on the framework gene *KIR3DL3* (assuming *KIR3DL3* always has two copies). When using PING, the thresholds of each CN group are manually determined. SakaueKIR uses average read depths directly calculated from the selected regions and clusters them by Kernel Density Estimation (KDE). Both PING and SakaueKIR require large sample sizes in a cohort for clustering. T1K estimates allele abundance by EM algorithm, but it limits to at most two alleles per gene, which does not meet the KIR complexity because a KIR gene may has more than two copies.

From allele typing perspective, the star alleles should be called with the estimated copy number of each gene. PING iteratively maps reads to the star allele candidates and gets the final 5-digit alleles. SakaueKIR employs a GATK-like pipeline and calls alleles by finding a set of star alleles that fits the partially phased SNPs calculated by HaplotyperCaller [12], but it gives ambiguous alleles because of the unreliable phased information. T1K tries to maximize the likelihood by choosing one or two alleles according to their abundance. However, T1K lacks the ability to call more than three alleles for a gene owing to its assumption. In summary, to the best of our knowledge, none of the existing methods can correctly type full-resolution KIR star alleles.

For clinical practices, it is better for a tool being able to update its KIR allele sequence database to keep up with the official KIR database Immuno Polymorphism Database-KIR [13] (IPD-KIR), which collects the names of star alleles and their corresponding sequences reported so far. Recent published tools like T1K can update its database, but SakaueKIR and PING are frozen to the version at which they are developed or released. SakaueKIR uses IPD-KIR 2.7.0 and PING uses IPD-KIR v2.7.1 stated in the published papers.

To conquer the difficulty oriented to the high similarity between KIR genes and KIR star alleles mentioned above, inspired by HISAT-genotype [14], we propose a novel pipeline, Graph-KIR. Similar to the solution provided by HISAT-genotype for typing HLA star alleles, Graph-KIR utilizes HISAT2, a graph read mapper, to map short reads to custom-built indexes. The highly accurate graph mapping enables Graph-KIR to estimate copy number per sample independently, thanks to the higher linearity between copy number and read depth in the graph alignment. The task of allele typing relies on the variants called from the graph, and a set of star alleles is called by maximum likelihood estimation. Furthermore, any version of IPD-KIR database can be converted to graph indexes using the tool provided by HISAT2 along with the custom codes together released with Graph-KIR.

In this study, Graph-KIR uses all the sequences in IPD-KIR v2.10.0, including 879 full-length sequences and 652 partial sequences (exon-only sequences). In total, 1,110 distinct star alleles are used in building the graph index. Each KIR gene has at least 30 alleles. When the performance is concerned, Graph-KIR underwent testing on both simulated and actual WGS samples. The annotation of actual samples had been previously conducted in another study.

## 2 Materials and Methods

### 2.1 Pipeline overview

Graph-KIR contains five main steps: (1) building graph index from MSA; (2) read mapping to the human genome and extracting KIR-related reads (optional); (3) graph read mapping and read filtering; (4) copy number estimation; and (5) allele typing, as illustrated in Figure 2.

**Figure 2.**
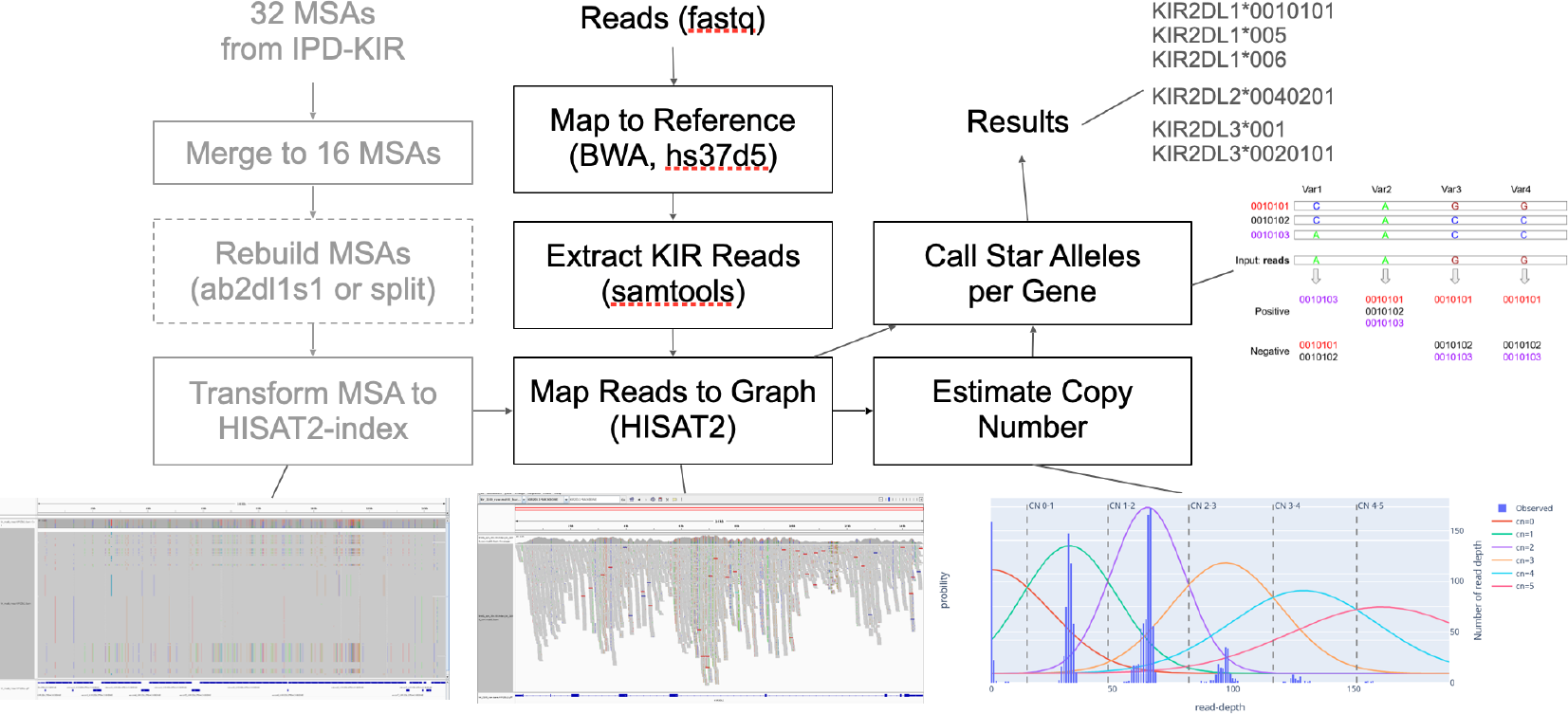
Overview of the Graph-KIR pipeline. The preprocessing steps that only need to be executed once are highlighted in grey, and the optional step is represented by a dashed box.

The graph index is built from the MSAs of all alleles for the 17 KIR genes available from IPD-KIR. These MSAs are manipulated by merging similar genes (*KIR2DL1* and *KIR2DS1*), left aligning, and adding pseudo introns to exon-only sequences, making the MSAs more suitable for graph index building and read mapping. Once the index is built, there is no need to rebuild it unless the database version of IPD-KIR changes.

The whole genome pair-end reads are first mapped to the human genome hg19. The reads mapped at KIR-related regions are extracted. If the input reads are already filtered for KIR regions, this step can be skipped.

Next, the reads are mapped to the graph index by HISAT2. Here, Graph-KIR discards multi-mapped reads and reads with low alignment scores because such reads may result in overestimation of read depth and cause false positives in variant calling.

The key step in copy number estimation of Graph-KIR is distribution fitting. Multiple distributions, each representing one of the copy numbers (0 to 6), are fit to read depths calculated from all the bases of each gene. The distributions have constraints about the linearity between copy number and the estimated read depth. This feature enables estimating copy numbers with only one sample while maintaining its robustness and high accuracy.

After copy number estimation, Graph-KIR calls a set of alleles for each gene with its size equal to the copy number determined previously. This allele typing step includes calling variants of reads from the graph, comparing the variants with the star alleles, assigning each read to an alleles in the set, and iteratively finding the best-fitting allele set.

### 2.2 MSA generation and index preparation

The MSAs are reconstructed from .msf and .dat files downloaded from IPD-KIR database. Each gene contains both exon-only and full-length MSA. In total, there are 32 MSAs from 17 genes, where *KIR2DL5A* and *KIR2DL5B* are already merged.

The MSAs in Graph-KIR are merged version that preserves both introns and exons information, combining the exon alignment from the exon-only MSA and the intron alignment from the full-length MSA. For exon-only alleles, i.e. sequences missing intronic regions, the introns are filled by consensus calculated from similar alleles in the same gene. The similar alleles are alleles that at least share the first 5 digits. If no allele are found, the first 3 digits are considered instead. In cases there is no similar alleles with at least the same first 3 digits, the consensus is calculated using all the full-length alleles in the same gene.

The recommended graph index for Graph-KIR index is ‘ab2dl1s1’, which has 15 MSAs. The *KIR2DL1* and *KIR2DS1* are merged and *KIR2DL5A* and *KIR2DL5* are merged. The other index for comparison is ab (16 MSAs, *KIR2DL5A* and *KIR2DL5B* are merged) and split (17 MSAs, no merged).

If it is required to merge two or more MSAs, Graph-KIR will split the MSA into blocks of MSAs by introns and exons. The MSA blocks correspond to the same intron or exon regions are realigned separately by muscle [15] with default parameters. Then, the merged MSA is constructed by concatenating the realigned blocks in order. Splitting the MSA, such as separating *KIR2DL5A* and *KIR2DL5B* from *KIR2DL5*, is a straightforward process. The split MSAs only include alleles with the corresponding gene name, and any column only contains gap is removed.

The last part of MSA processing is left-aligning, inspired by the VCF normalization [16]. Here, all deletions in the MSA will be left-aligned.

Once MSAs are prepared, the consensus can be calculated by selecting the base with maximum frequency in each column while ignoring gaps. These consensus sequences are then saved in the .fasta format as the graph backbone. The SNPs, called from each star allele against the backbone, are saved in .snp and .haplotype files. Note that SNPs with frequency greater than 0.1 will be reserved in the graph index. Graph indexes are then built by running ‘hisat-build’ with the backbone and SNPs files mentioned above.

The MSA operations previously described are from another open-source project developed by the same authors of Graph-KIR. The codes are available at https://github.com/linnil1/pyHLAMSA.

### 2.3 Read Extraction

Before entering to core Graph-KIR algorithm, the WGS (whole genome sequencing) reads are first mapped to hg19 reference. Graph-KIR uses the version hs37d5 of hg19 obtained from 1000 Genomes Project [17]. Read mapping is run by BWA-MEM [18] with parameters -M -K 100000000 -a. Then, the reads mapped to KIR regions are extracted via Samtools [19] but exclude singleton. The KIR regions are 19:55200000 to 19:55400000 and GL000209.1, or chr19:55200000-55400000 and chr19_gl000209_random if UCSC [20] version is used. Currently, we use hg19 instead of hg38 because hg38 contains numerous KIR-related scaffolds that need cautious examination.

### 2.4 Read mapping

The read mapping parameters we used are similar to HLA typing in HISAT2-genotype `hisat2 --no-spliced-alignment --max-altstried 64 –haplotype`. In this research, the extracted reads are mapped to the three different graph indexes: ‘ab2dl1s1’, ‘ab’, and ‘split’ for the following comparison.

After mapping, we remove reads with multiple mapping, low alignment score (edit distance > 4) or unpaired flag. Those discarded reads mostly came from other KIR genes.

### 2.5 Copy number estimation

The copy number estimator is a composite of multiple distributions. Each distribution represents the read depths of genes with identical copy number. Thus, the distributions correspond with different non-negative integer copy number. Furthermore, the parameters of distributions exhibit a linear relationship with the corresponding copy number. These characteristics rely on the assumption that the reads are uniformly sequenced across the genome and be accurately mapped by the read mapper to their respective genes.

The read depth of a gene is determined as the 75th percentile of the read depths across the gene. There are alternatives such as average or median. If WES/capture-based samples are provided, Graph-KIR has the option to calculate the depths with the exon part only.

Let *D* = {*d*_2DL1_, *d*_2DL2_,…, *d*_*g*_} be the read depth of each gene. Here, we assume the read depths of genes with copy number k follow a normal distribution 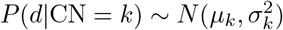. The parameters of distributions are constrained to *μ*_*k*_ = *k*. *μ*_2_/2 because of the linear relationship mentioned above. Additionally, the standard deviations are adjusted so that read depth is more likely to fit to CN=2 instead of other copy numbers. Here, the deviations at CN=2 are kept minimal, while the other deviations are increased to reduce their likelihood. Furthermore, CN=1 is more likely to occur than CN=3. To satisfy this preference, we introduce two constants, *C*_*n*_ and *C*_*p*_, where *C*_*n*_ is smaller than *C*_*p*_. This allows the equation to take the following form:

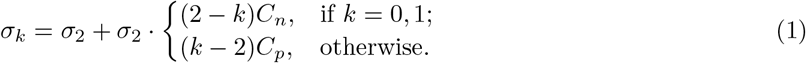

Once *μ*_2_ and *σ*_2_ are set, the distributions are all fixed. The copy number of a gene can be determined by assigning it to the distribution CN=*k* that has the maximum likelihood of observing its read depth:

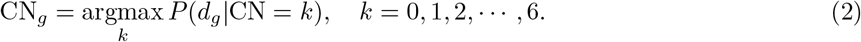

The overall likelihood of observing the inferred copy numbers given the distributions can be calculated by multiplying the likelihood of each gene described in the equation (2). The maximum can be found by changing the variable *μ*_2_, the others are set to *σ*_2_ = 0.08, *C*_*p*_ = 0.5, *C*_*n*_ = 0.3 and read depths are normalized by the maximum one in this study.

In rare cases, for example, all of the genes are CN=1 and CN=2, or all of the genes are CN=2 and CN=4. Those two cases are ambiguous for Graph-KIR maximum-likelihood fitting. The method will tend to predict CN=1 and CN=2 because *C*_*n*_ *< C*_*p*_.

Graph-KIR provides an option to assume *KIR3DL3*, one of the framework genes, has two copies and adjust *μ*_2_ based on the assumption. Additionally, Graph-KIR offers an additional option to determine the copy number of multiple samples jointly. The only change in this option is that the read depths are calculated from all the samples in the cohort, i.e. 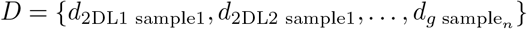.

### 2.6 Allele typing

The allele typing is, indeed, finding a possible set of star alleles according to the variants in the reads. The number of alleles (set size) of each gene is the copy number estimated in the previous step. Graph-KIR determined the set of each gene independently.

In the first step of allele typing, Graph-KIR calculates the likelihood of the read originating from an specific allele by matching the variants within the read interval. For any unmatched variant, a probability of 0.001 is assigned, indicating the probability of sequencing errors or mapping errors. The probability is defined as

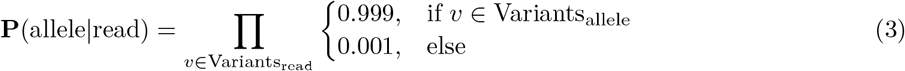

where Variants_allele_ is variants inside the allele and Variants_read_ is variants inside the read.

Subsequently, the overall probability of the reads being generated from a set of alleles (denoted as ‘alleles’) is calculating the product of assigning each read to one of the alleles that has the maximum probability:

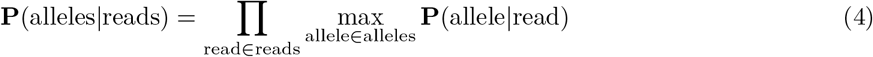

In the final step, the alleles for each gene can be determined by choosing the set that has the highest overall probability from all possible combinations:

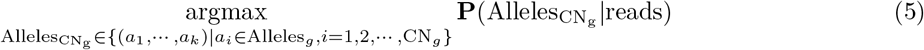

where ‘Alleles_*g*_’ is all the alleles of a gene *g*, and 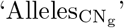 is the determined allele set of a gene.

Instead of examining all combinations which is computationally intensive, our strategy is to iteratively increase alleles set size from one until reaching the estimated copy number. For each iteration, we only preserve the top 600 sets having highest overall probability.

To minimize the error caused by mapping error and sequencing error, Graph-KIR does some error correction treatment, which ignores a variant if it has read depth less than 3 or the variant is rare (The read depth ratio of alternative over reference type < 0.25).

### 2.7 Simulated data preparation

To evaluate whether our entire workflow works correctly, simulated data is used for testing before running on real-world data. The same pipeline as PING [8] is used to generate simulated short reads. In short, two haplotypes are randomly selected from the haplotypes summarized by [21] for a human sample. Once the haplotypes are chosen, copy number for each KIR gene can be collected, and then KIR alleles correspond to each gene are randomly selected with replacement. After all the alleles are selected, ART [22] will generate reads from the sequences of selected KIR alleles. In total, 100 samples are generated and all of them contain about 30x 150bp paired-end reads with sequencing errors. Note that the sequences generated by this method don’t include the inter-gene sequences because this research mainly focuses on KIR gene content.

### 2.8 HPRC data

The real sample set for evaluating the performance of KIR pipelines are from Human Pangenome Reference Consortium (HPRC) [23]. In total, 44 out of 47 samples have WGS short pair-end reads available from NCBI. Here is the list of HPRC samples: HG002, HG00438, HG005, HG00621, HG00673, HG00733, HG00735, HG00741, HG01071, HG01106, HG01109, HG01175, HG01243, HG01258, HG01358, HG01361, HG01891, HG01928, HG01952, HG01978, HG02055, HG02080, HG02109, HG02145, HG02148, HG02257, HG02572, HG02622, HG02630, HG02717, HG02723, HG02818, HG02886, HG03098, HG03453, HG03486, HG03492, HG03516, HG03540, HG03579, NA18906, NA19240, NA20129, NA21309.

The annotation of HPRC samples used in this research was obtained in a previous study [24]. The annotated KIR alleles were identified by using minimap2 [25], where allele sequences from the IPD-KIR database were mapped to high-quality phased haplotypes, which were generated by assembling HiFi reads from each HPRC sample. The annotated alleles were further validated by manual examination using IGV [26]. Notably, numerous novel alleles were found, but for subsequent comparisons, the closest alleles were considered.

### 2.9 Allele comparison

To evaluate the result of allele typing, we compare the predicted alleles with the ground truth alleles at four different resolutions: 7-digit, 5-digit, 3-digit, 0-digit (gene level). A pair of predicted allele and ground truth allele will be marked as correct if they share exactly the same or more digits at the desired level. Additionally, each predicted allele and ground truth allele can only be paired with at most one of its counterpart. Accuracy at each resolution is calculated as the number of correct pairs divided by the number of ground truth allele. Unpaired predicted alleles and answer alleles will be marked as False Positive (FP) and False negative (FN), respectively.

If a tool outputs multiple sets of alleles when it encounters ambiguity (PING, SakaueKIR), we simply choose the first set to evaluate. It ensures that the number of alleles is equal to its estimated copy number. Another evaluation is to treat all output sets as the called alleles for a sample. This treatment will create lots of False Positive (FP) alleles, but the accuracy will be higher because one of the predicted alleles may match the ground truth allele. In the case of novel alleles, the alleles provided by the tool will be used for comparison.

## 3 Results

### 3.1 Read mapping

The effectiveness of Graph-KIR heavily relies on the precision of graph read mapping. Improvements in read mapping can subsequently enhance the results of both copy number estimation and allele typing. To assess the effectiveness of read mapping, especially when multiple mapping for a read is possible, we utilize metrics including False Discovery Rate (FDR), recall, precision, and alignment rate. FDR represents the proportion of incorrect mapping positions of all the reads including multiple mapping. Recall is the rate of reads that have at least one correct mapping position. Precision is 1 - FDR. The alignment rate quantifies the proportion of mapped reads among all the reads of each gene.

Various methods were assessed here, including linear mapping (BWA-MEM, Bowtie 2) and graph mapping (Graph-KIR) with different indexes (‘split’, ‘ab’, ‘ab2dl1s1’). This evaluation is conducted on the 100 simulated samples. All setups with the ‘unique’ option (points in cross) have a smaller recall but also a smaller FDR compared to the ‘all’ option (points in circle) shown in Figure 3. This phenomenon indicates that filtering multiple mapped reads leads to a decreased numerator in recall, but it simultaneously decreases the number of false positives, resulting in smaller numerator and denominator in FDR. Overall, recall rates slightly dropped while FDRs significantly decreased.

**Figure 3.**
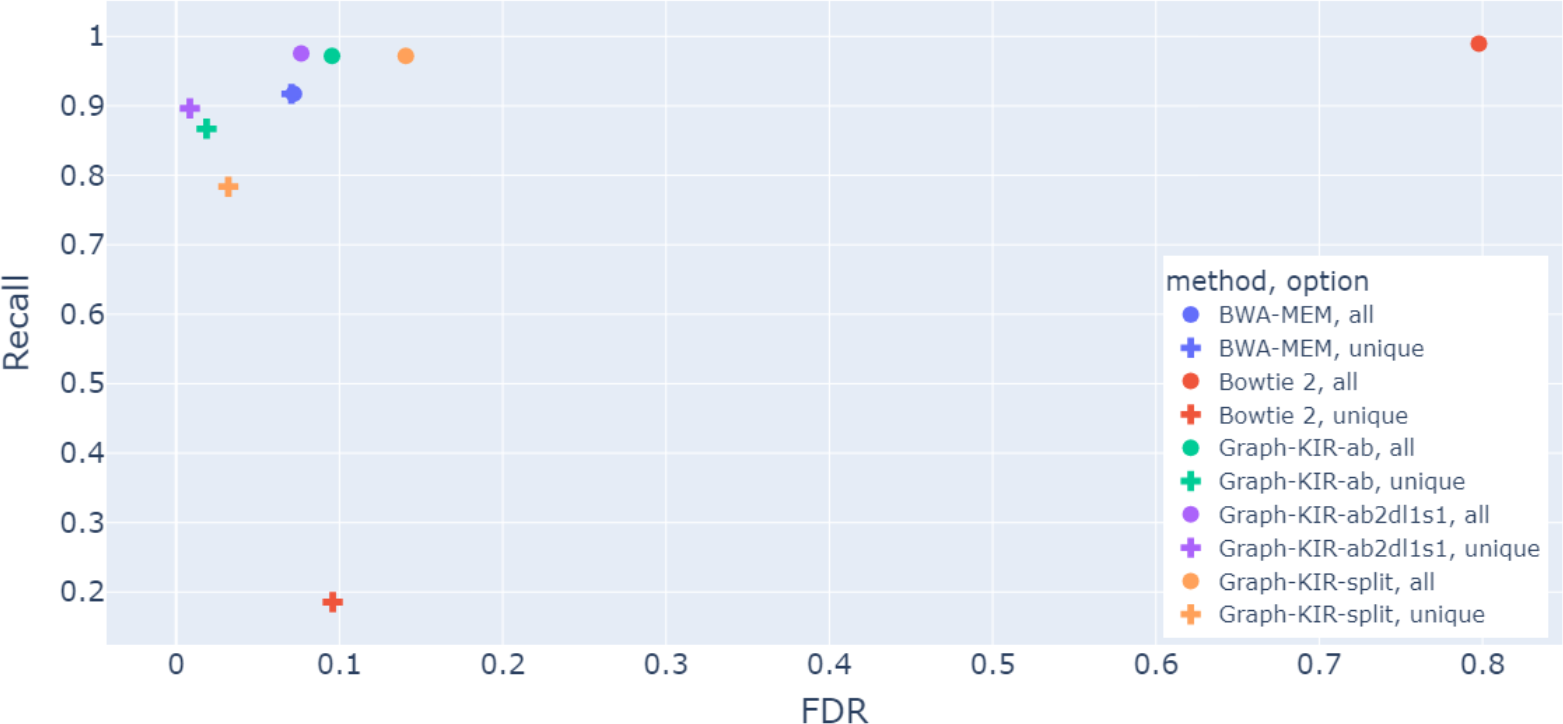
The effectiveness of various mapping strategies is assessed based on False Discovery Rate (FDR) and Recall of the reads across 100 simulated samples. Recall is the ratio of the reads having at least one correct mapping position divided by the total number of reads. Conversely, FDR is the percentage of mapping positions that are incorrect. For the methods with the ‘unique’ suffix, multi-mapped reads are disregarded, while ‘all’ indicates all the reads are considered. Both BWA-MEM and Bowtie 2 use ‘ab2dl1s1’s backbone as the index.

In Figure 3, Bowtie 2 has the highest false discovery rate (FDR = 0.81). Among the three different graph indexes, Graph-KIR based on ‘split’ (17 graphs) has the highest FDR (FDR = 0.20). Index ‘ab’ (16 graphs) merges two similar genes *KIR2DL5A* and *KIR2DLB* to reduce FDR to 0.12. Index ‘ab2dl1s1’ further lowers the FDR to 0.09 by merging *KIR2DL1* and *KIR2DS1*. These three results show that merging similar genes to eliminate sequence redundancy and allele similarity between indexes can decrease the FDR of the graph mapper. The index ‘ab2dl1s1’ of Graph-KIR is a little less accurate than BWA-MEM (‘BWA-MEM’, FDR=0.07). However, the unique-filtering version ‘ab2dl1s1, unique’ decreases FDR (0.09 to 0.008) at the expense of a lower recall rate (0.97 to 0.89). Applying the same filtering on BWA-MEM (‘BWA-MEM, unique’) does not result in an apparent decrease in the FDR (0.071 to 0.069).

To compare the read mapping results of different strategies thoroughly, we also take precision and alignment rate at gene-level into account, as shown in Figure 4. In this figure, all methods have been filtered using the ‘unique’ option. Each method has 17 points, representing 17 KIR genes. The results demonstrate that the proposed method, Graph-KIR with the ‘ab2dl1s1’ index, has significantly higher precision than other methods. Although BWA-MEM exhibits higher alignment rate, which means preserving more reads, some of the retained reads may include noise at downstream allele typing due to the lower precision. In conclusion, Graph-KIR outperforms other linear mapping methods in terms of precision, FDR, and recall rate. Among all the indexes evaluated, ‘ab2dl1s1’ stands out as the most suitable one.

**Figure 4.**
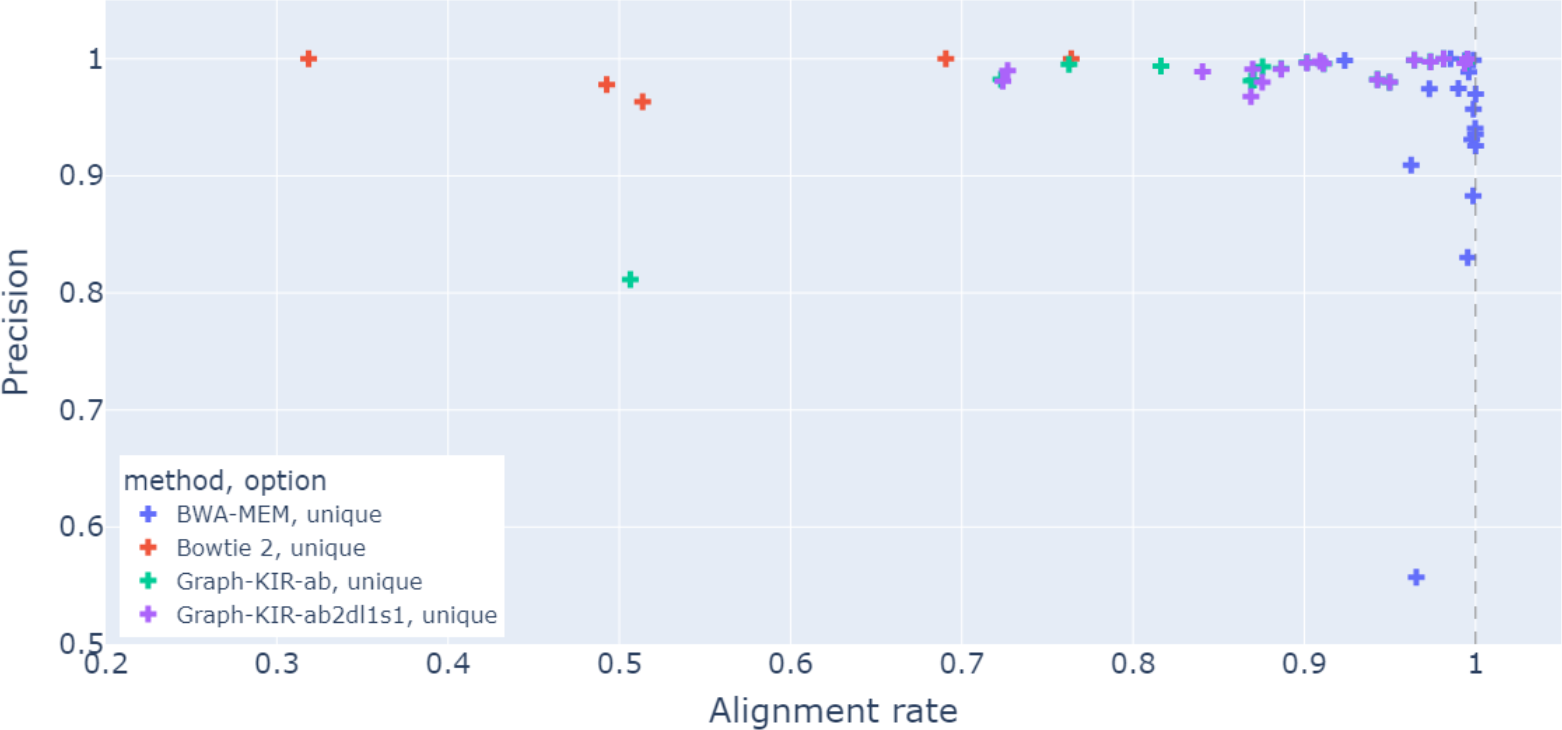
Precision is plotted against alignment rate for each method at gene level, with 17 points per method. Precision is the proportion of reads that are correctly mapped to their originating locations. The alignment rate quantifies the proportion of mapped reads among all the reads of each gene. For the methods with the ‘unique’ suffix, multi-mapped reads are disregarded. Both BWA-MEM and Bowtie 2 use ‘ab2dl1s1’s backbone as the index.

### 3.2 Copy number estimation

The performance of four other KIR pipelines and different options of Graph-KIR on copy number estimation was evaluated by 100 simulated data, which has ground truth as it was generated. The difference against the ground truth is calculated by the sum of the absolute difference of each gene across all simulated samples, denoted as ‘Diff’ in Table 1.

**Table 1:** Performance of copy number estimation on 100 simulated samples. Two stages of copy number estimation (‘Before allele typing’ and ‘After allele typing’) are performed. The first stage directly utilizes the estimated copy numbers while the second stage obtain the copy numbers from the allele typing results. The first stage of PING’s copy number result (‘Before allele typing’) is obtained from ‘manualCopyNumberFrame.csv’, which is generated after manually providing the copy number thresholds. KIR*KPI only indicates the presence/absence of a gene but does not provide copy number information. Therefore, we use the predicted haplotypes to infer the copy number of each gene (by ‘haps.txt’).

| Methods | Total | Before allele typing |  | After allele typing |  |
| --- | --- | --- | --- | --- | --- |
|  |  | Diff | Accuracy | Diff | Accuracy |
| Graph-KIR-split | 2389 | 256 | 0.892 | 256 | 0.892 |
| Graph-KIR-ab | 2389 | 95 | 0.960 | 101 | 0.957 |
| Graph-KIR-ab2dl1s1 | 2389 | 0 | 1.000 | 20 | 0.991 |
| Graph-KIR-ab2dl1s1-cohort | 2389 | 0 | 1.000 | 20 | 0.991 |
| Graph-KIR-ab2dl1s1-median | 2389 | 35 | 0.985 | 53 | 0.977 |
| Graph-KIR-ab2dl1s1-exon | 2389 | 11 | 0.995 | 31 | 0.987 |
| KIR*KPI | 2389 | 389 | 0.837 | 389 | 0.837 |
| PING-wgs | 2389 | 2 | 0.999 | 30 | 0.987 |
| SakaueKIR | 2389 | 627 | 0.737 | 1325 | 0.445 |
| T1K | 2389 | NA | NA | 321 | 0.865 |

‘split’ indicates that the graph index contains 17 graphs, where no gene is merged.
‘ab’ indicates that the graph index contains 16 graphs, where *KIR2DL5A* are *KIR2DLB* merged.
‘ab2dl1s1’ indicates that the graph index contains 15 graphs, where *KIR2DL5A* and *KIR2DLB* are merged, and *KIR2DL1* and *KIR2DS1* are merged.
‘cohort’ indicates that copy numbers are jointly called rather than independently from each sample.
‘median’ indicates that the median of read depths is used instead of the 75th percentile.
‘exon’ indicates that only the read depth of the exon part is used.
‘wgs’ indicates the wgs version of PING is used for comparison.
Accuracy = 1 – Diff/Total.
NA: not available.

However, most KIR typing pipelines treat the copy number as an intermediate result, so they do not always separately output the copy number for each gene. For instance, some pipelines aggregate the copy numbers of *KIR2DL5A* and *KIR2DL5B* and assign the combined value to a new gene named *KIR2DL5*. To calculate the difference of these merged genes against ground truth, the evaluation takes this into account and compares the copy number with the total value derived from summing up copy numbers of the corresponding genes. Another measurement is to derive copy numbers from allele typing results instead of intermediate copy numbers. It may potentially be a more unbiased evaluation for the merged genes.

As summarized in Table 1, among the methods of Graph-KIR with three indexes, ‘ab2dl1s1’ (default) has 100% accuracy, ‘ab’ has much lower accuracy, and ‘split’ has the lowest accuracy. We conclude that merging similar KIR genes preserved more uniquely mapped reads for read depth calculation, which highly influences the copy number estimation results. Among different configurations, three key points can be concluded:

- Using median value (‘-median’) to represent the read depth of a gene is not better than using the 75 percentile (default).
- Determining the copy number by cohort (‘-cohort’) and one-by-one version (default) perform equally well.
- Calculating read depth using only the exon region (‘-exon’) has an accuracy lower than using full sequence (default).

Among the four methods for comparison, PING[9] has almost perfect copy number prediction when the correct threshold was manually applied. We manually set the threshold by comparing the value against the ground truth of copy numbers. Other tools like KIR*KPI, SakaueKIR, and T1K all have accuracy below 0.86.

Table 2 presents the accuracy of estimated copy numbers across 44 HPRC samples. The results are similar to Table 1, with Graph-KIR outperforming the other tools. The superior performance of Graph-KIR’s per-sample estimation, compared to cohort estimation, can be attributed to the varying sequence depths across some of the HPRC samples, i.e., not always 30x. Graph-KIR does not normalize read depth of genes using the depth of framework genes (e.g., *KIR3DL3*), as such normalization is not always flawless given that *KIR3DL3* may not be diploid in rare instances.

**Table 2:** Performance of copy number estimation on 44 HPRC samples. Two stages of copy number estimation (‘Before allele typing’ and ‘After allele typing’) are performed. The first stage directly utilizes the estimated copy numbers while the second stage obtain the copy numbers from the allele typing results.

| Methods | Total | Before allele typing |  | After allele typing |  |
| --- | --- | --- | --- | --- | --- |
|  |  | Diff | Accuracy | Diff | Accuracy |
| Graph-KIR-ab2dl1s1 | 832 | 4 | 0.995 | 34 | 0.959 |
| Graph-KIR-ab2dl1s1-cohort | 832 | 17 | 0.979 | 45 | 0.945 |
| PING-wgs | 832 | 65 | 0.921 | 106 | 0.872 |

‘ab2dl1s1’ indicates that the graph index contains 15 graphs, where *KIR2DL5A* and *KIR2DLB* are merged, and *KIR2DL1* and *KIR2DS1* are merged.
‘cohort’ indicates that copy numbers are jointly called rather than independently from each sample.
‘wgs’ indicates the wgs version of PING is used for comparison.
Accuracy = $1 - \text{Diff}/\text{Total}$ .

### 3.3 Allele typing

The performance of allele typing using Graph-KIR with different configurations and other KIR pipelines across the 100 simulated samples are shown in Table 3. The results are presented at 4 resolutions: 7-digit, 5-digit, 3-digit, and gene-level. The index ‘ab’ is slightly inaccurate (about 3%) than ‘ab2dl1s1’ at all resolutions, probably due to the error in copy number estimation as described in previous section, and it can be further validated by the low accuracy in typing *KIR2DS1* demonstrated in Figure S1.

**Table 3:** Performance of allele typing on 100 simulated samples. The numbers in the parentheses stand for the total count of alleles at that resolution.

| Correct count | 7-digit<br>(1752) | 5-digit<br>(1946) | 3-digit<br>(2389) | Gene<br>(2389) |  |  |
| --- | --- | --- | --- | --- | --- | --- |
| Graph-KIR-ab2dl1s1 | 1597 | 1888 | 2321 | 2379 |  |  |
| Graph-KIR-ab | 1551 | 1836 | 2249 | 2291 |  |  |
| PING-wgs-ans | NA | 1789 | 2244 | 2369 |  |  |
| SakaueKIR | 107 | 865 | 1069 | 1591 |  |  |
| SakaueKIR-allposs | 528 | 1110 | 1367 | 1819 |  |  |
| SakaueKIR-ans | 120 | 991 | 1223 | 1615 |  |  |
| SakaueKIR-ans-allposs | 513 | 1087 | 1352 | 1812 |  |  |
| T1K | 411 | 1646 | 2051 | 2082 |  |  |
| Accuracy | 7-digit | 5-digit | 3-digit | Gene | FN | FP |
| Graph-KIR-ab2dl1s1 | 0.911 | 0.970 | 0.971 | 0.995 | 10 | 10 |
| Graph-KIR-ab | 0.885 | 0.943 | 0.941 | 0.958 | 98 | 3 |
| PING-wgs-ans | NA | 0.919 | 0.939 | 0.991 | 20 | 10 |
| SakaueKIR | 0.061 | 0.444 | 0.447 | 0.665 | 798 | 585 |
| SakaueKIR-allposs | 0.301 | 0.570 | 0.572 | 0.761 | 570 | 51565 |
| SakaueKIR-ans | 0.068 | 0.509 | 0.511 | 0.676 | 774 | 551 |
| SakaueKIR-ans-allposs | 0.292 | 0.558 | 0.565 | 0.758 | 577 | 24653 |
| T1K | 0.234 | 0.845 | 0.858 | 0.871 | 307 | 14 |

‘ab’ indicates that the graph index contains 16 graphs, where *KIR2DL5A* are *KIR2DLB* merged.
‘ab2dl1s1’ indicates that the graph index contains 15 graphs, where *KIR2DL5A* and *KIR2DLB* are merged, and *KIR2DL1* and *KIR2DS1* are merged.
‘allposs’ indicates all allele sets in output of a tool are treated as called alleles; otherwise, only the first set is used.
‘ans’ indicates the copy number is assigned by peeking the ground truth.
‘wgs’ indicates the wgs version of PING is used for comparison.
Graph-KIR and T1K are both based on IPD-KIR version 2.10.0. PING-wgs is based on IPD-KIR 2.11.0. SakaueKIR is based on 2.7.0
False positive (FP) indicates the number of unpaired predicted alleles while false negative (FN) indicates the number of unpaired answer alleles.
NA: not available.

Graph-KIR once again demonstrates higher accuracy than other KIR tools, which shows 0.87 and 0.91 at 7-digit resolution and 0.93 and 0.97 at 5-digit resolution. PING has 0.92 5-digit accuracy but lacks the ability to provide 7-digit resolution. T1K has 0.84 accuracy at 5-digit resolution and very low accuracy (0.24) at 7-digit resolution. The false negative of T1K is quite large because it predicts at most 2 alleles per gene which is not always true for KIR genes. SakaueKIR has the lowest accuracy (0.57) at 5-digit resolution, mainly because most of the alleles in simulated data are novel for the IPD version 2.7.0. Since there’s no command for updating its database, this is the best results we can achieve for it.

Table 4 shows the allele typing results compared to the annotated HPRC samples. Only PING and Graph-KIR are selected for comparison because of their superior performance on the simulated data. Notably, numerous alleles are found to be novel alleles in IPD-KIR version 2.12.0 or having fusion with another gene, hindering the HPRC dataset to benchmark performance of the tools. We manage to retain the alleles with high confidence annotation and the results showcase that Graph-KIR remains superior to the alternative regarding of accuracy at all resolutions while maintaining a low number of false negatives.

**Table 4:** Performance of allele typing on 44 HPRC sample. The column ‘Filter’ stands for the types of alleles removed from the evaluation.

| Filter | Methods | 7-digit | 5-digit | 3-digit | Gene | FN | FP |
| --- | --- | --- | --- | --- | --- | --- | --- |
| -fusion | number of alleles | 248 | 717 | 827 | 827 |  |  |
|  | Graph-KIR | 0.706 | 0.736 | 0.747 | 0.979 | 17 | 17 |
|  | PING-wgs | NA | 0.647 | 0.653 | 0.878 | 101 | 13 |
| -fusion -# | number of alleles | 237 | 654 | 741 | 741 |  |  |
|  | Graph-KIR | 0.713 | 0.743 | 0.765 | 0.978 | 16 | 17 |
|  | PING-wgs | NA | 0.702 | 0.718 | 0.881 | 88 | 12 |
| -fusion -# -+ -= | number of alleles | 222 | 520 | 552 | 552 |  |  |
|  | Graph-KIR | 0.721 | 0.813 | 0.864 | 0.978 | 12 | 17 |
|  | PING-wgs | NA | 0.715 | 0.739 | 0.882 | 65 | 12 |
| -fusion -# -+ -= -\$ | number of alleles | 146 | 147 | 149 | 149 | | |
|  | Graph-KIR | 0.705 | 0.850 | 0.893 | 0.980 | 3 | 15 |
|  | PING-wgs | NA | 0.755 | 0.785 | 0.893 | 16 | 10 |

#: an allele having nonsynonymous variant(s) in CDS, segmental deletion or fusion with another gene

## 4 Discussion

### 4.1 Similarity between KIR Genes

Genes with high similarity often lead to reads being mapped to multiple positions, introducing ambiguity and complicating subsequent analyses. We attempted to address this challenge by merging similar genes before building graph index for read mapping, such as *KIR2DL5A* and *KIR2DL5B*, resulting in improved performance in both steps.

For Graph-KIR, merging *KIR2DL1* and *KIR2DS1* is critical, especially for copy number estimation. Many reads are mapped to both *KIR2DS1* and *KIR2DL1* in the ‘ab’ index. Only half (51%) of the reads mapped to *KIR2DS1* and 87% of the reads mapped to *KIR2DL1* are uniquely mapped. Removing the multi-mapped reads significantly influenced the read depth of these genes, leading to errors in copy number estimation. In contrast, in the ‘ab2dl1s1’ index, 87% of *KIR2DL1* reads and 84% of *KIR2DS1* reads are retained.

### 4.2 Limitation

Graph-KIR does not consider the phasing information across different pairs of reads. This means that when the distance between variants exceeds the short read length, the algorithm is unable to determine the correct allele set which variants belong to because they all share the same likelihood. Consequently, this limitation may result in incorrect allele typing for a different set of alleles. The possible sets of alleles with identical likelihood are saved in a separate file in the Graph-KIR pipeline.

To measure the influence of the limitation mentioned above, we compare the potential allele sets and the ground truth alleles. By choosing the allele set that matches the ground truth alleles the most for each gene, we discover the accuracy in allele typing increase from 0.91 to 0.98. This suggests that approximately 6% of the error is owing to this limitation.

While multi-mapped reads are currently discarded after the Graph-KIR read mapping steps to enhance precision, distinguishing them from multiple mapping positions could lead to improved performance in both precision and recall rate.

During allele typing, Graph-KIR doesn’t favor common alleles, i.e. all alleles are treated equally. Some tools [27] might implement this concept to achieve a higher accuracy. However, in this study, we assert that KIR alleles are still underestimated, suggesting that prior knowledge of allele frequency might not be particularly beneficial. Thus, the favor of a common allele is not implemented for Graph-KIR’s allele typing.

### 4.3 Evaluation of Speed

Graph-KIR uses existing tools for computation-intensive tasks and only customizes the core function of copy number estimation and allele typing. For example, BWA-MEM for read mapping and extraction, HISAT2 for graph mapping and Samtools for read depth calculation. These optimized tools speed up the execution time of Graph-KIR. In the test of 100 simulated samples, it takes Graph-KIR 41 minutes to sequentially process each sample, and takes 8 minutes if run in parallel e.g. GNU Parallel [28] because Graph-KIR can predict KIR allele per sample independently. In contrast, it takes PING about 52 hours 20 minutes for the 100 samples, including the time for the in-house program to automatically provide the thresholding information. All the experiments were conducted on high-performance computer TAIWANIA3’s SLURM partition ngs92G, which has 14 virtual cores and 92G virtual memory.

## 5 Conclusion

Graph-KIR is a brand new pipeline for typing KIR alleles at 7-digit resolution from short-read WGS data. With a proper index and configurations selected, the polymorphism of KIR genes can be effectively handled by the tool optimized for graph mapping. The procedure to determine copy numbers can be done with only one sample, this feature enables KIR research without collecting lots of samples. When evaluated with both simulated and real data, the superior performance of Graph-KIR over other existing KIR tools (T1K and SakaueKIR) is obvious. While PING shows competitive performance with Graph-KIR at lower resolution, it has the disadvantage of requiring manual thresholding and also suffers from longer runtime.

Currently, Graph-KIR only accepts short-read data. A feature that supports analysis of long-read data to complement the phasing problem encountered with short-read data will be added to a future version. In addition, the core algorithm to determine copy numbers is heavily impacted when most KIR genes have the same copy number. Refining the algorithm by estimating the variable *μ*_2_ from other genes reported to be generally having two copies can be considered. Also, the pipeline can be modified and adapted for HLA allele typing, especially for genes have copy number issue like *DRB3, DRB4* and *DRB5*[29, 30].

## Code availability

Graph-KIR and research-related codes are available at https://github.com/linnil1/KIR_graph.

### Acknowledgements

We thank National Center for High-performance Computing (NCHC) for providing computational and storage resources.

