## Supplemental file 1 for "Graph-KIR: Graph-based KIR Copy Number Estimation and Allele Calling Using Short-read Sequencing Data"

### Supplement

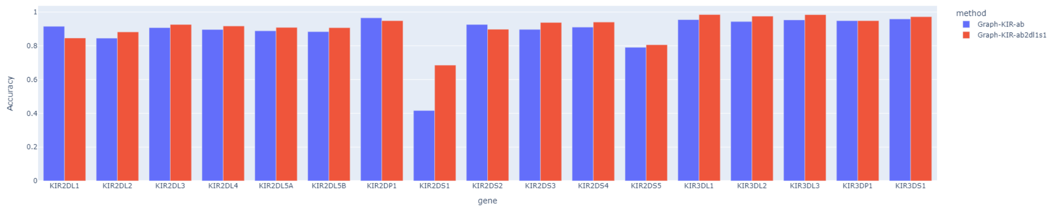


Figure S1: The accuracy of allele typing at 7-digit resolution on the simulated data. Each KIR gene is evaluated separately.
